# BfBio: a graph-based tool for the prediction of Angiogenic Stalk Cell genes using a Personalized PageRank algorithm

**DOI:** 10.64898/2026.08.23.746493

**Authors:** Léo Bettoni, Julia Dmitrieva, Mira Mousa, Habiba Alsafar, Yvan Saeys, Pooya Zakeri, Nuphar Veiga, Peter Carmeliet

**Affiliations:** Department of Oncology, Laboratory of Angiogenesis and Vascular Metabolism, KU Leuven, Leuven, Belgium; Laboratory of Angiogenesis and Vascular Metabolism, VIB Center for Cancer Biology, Leuven, Belgium; Center for Biotechnology, Khalifa University for Science and Technology, Abu Dhabi, UAE; Department of Public Health and Epidemiology, College of Medicine and Health Sciences, Khalifa University for Science and Technology, Abu Dhabi, UAE; Department of Biomedical Engineering and Biotechnology, College of Medicine and Health Sciences, Khalifa University for Science and Technology, Abu Dhabi, UAE; Data mining and Modelling for Biomedicine, VIB-UGent Center for Inflammation Research, Ghent, Belgium; Scientific Computing and Machine Learning, Department of Informatics, Faculty of Mathematics and Natural Sciences, University of Oslo, Oslo, Norway

## Abstract

Although most human protein coding genes have functional annotations in databases, such as GeneCards, many remain poorly characterized. To address this gap, computational tools can be leveraged to predict the functional roles of under-annotated genes by extracting patterns from complex biological networks.

Here we introduce Brain-for-Biotech (BfBio), a framework designed to identify genes important for vascular endothelial cells (EC), which are crucial cells for vessel formation (angiogenesis), vascular homeostasis, hemostasis and blood/tissue barrier function but also critical mediators of immunity and cancer progression. BfBio utilizes a Personalized PageRank (PPR) algorithm on an integrated network of different omics datasets and publicly available gene-gene/protein-protein interaction databases. In this study, we apply BfBio’s predictive capabilities to infer angiogenic stalk cell phenotype function in genes for which this function was not known before.

By leveraging a set of genes characterizing the stalk cell cluster in lung tumor EC models previously identified, we have achieved a high Area Under Receiver Operative Characteristic (AUC-ROC) performance (0.837). Enrichment analysis, coupled with a text mining application, further confirmed that among the 49 predicted genes four of them were poorly characterized yet possessed biologically relevant properties and were linked to cancer, thereby validating BfBio as a robust tool for prioritizing novel therapeutic targets in vascular biology.

**Author summary:** A third of the human coding genome remains poorly annotated, representing a potential goldmine for target discovery and drug development. There is a daunting and constant need for new drugs, especially for angiogenesis, which relates to the formation of new blood vessels from pre-existing ones made of endothelial cells. One specific subtype called stalk cells play a key role in cancer progression, promoting the tumor hypervascularization. Therefore, identifying genes responsible for this stalk cell phenotype has a significant therapeutic importance.

After creating a biological integrated network from protein-protein interaction databases and endothelial stalk cell-specific omics datasets, we have combined the use of gene prioritization, relying on a Personalized PageRank algorithm, with text mining to identify poorly annotated genes with a potential endothelial stalk cell phenotype function. We have prioritized four promising potential novel targets according to preliminary druggability assessment. These genes represent prime candidates for further experimental characterization to elucidate their role in angiogenesis and the tumor microenvironment, which is already hinted by their up-regulation in several cancer-related diseases.

## Introduction

Angiogenesis is a biological process that involves the formation of new blood vessels from pre-existing ones, which are made among others of endothelial cells (EC). Upon stimulation by hypoxia or growth factors, EC can switch from a quiescent state to active growth through either tip or stalk cell differentiation [1], two EC subtypes that are respectively responsible for leading the vascular sprouting by navigating at the vascular forefront and elongating the newly-generated branch following this direction. Within the tumor microenvironment, EC undergo a profound functional reprogramming to adopt a pro-angiogenic phenotype, characterized by proliferation, migration and sprouting [2, 3]. By forming a new network of blood vessels, sprouting angiogenesis serves as a critical tipping point in cancer progression, supporting the metabolic demands and rapid growth of the tumor mass [4]. Due to their crucial role in vascular sprouting, stalk cells are therefore key drivers of tumor hypervascularization [5]. As a result, identifying genes responsible for this phenotype has considerable significant therapeutic importance [6]. Yet, many regulators of the tumors EC (TEC) function remain poorly characterized, highlighting the need for the development of systematic approaches to uncover novel drivers of pathological vascular transitions.

Originally designed to rank Internet pages according to their importance to a specific web search [7], the PageRank (PR) algorithm provides a robust mathematical framework for prioritizing nodes based on their global connectivity within a network. More specifically, in this model the importance of a specific node is determined by the average importance of its neighbors having incoming links to it [8]. Since complex biological systems comprise thousands of interacting entities, often represented as multi-layered networks of pathways, protein-protein interactions, disease networks and drug-gene/disease-gene association, the PR algorithm offers powerful means to prioritize key functional drivers within their broader molecular environment. By leveraging the inherent structure of these biological networks, PR-based approaches can identify central nodes that govern systemic responses, providing a systematic way to rank genes within their broader molecular context.

In this work, we utilized a variant of the original PR algorithm, called Personalized PageRank (PPR) designed to rank nodes in a network based on their connectivity to specific nodes of interest, to elaborate on the inference of angiogenic stalk cells properties. We constructed networks from multiple EC bulk transcriptomic datasets using Pearson’s correlation and systematically optimized the model by exploring different combinations of correlation coefficient *r* and damping factor *α*. Recognizing that integrating multiple modalities capturing different aspects of the data enhances the predictive power, we augmented co-expression information from transcriptomic datasets with protein-protein interaction (PPI) networks. Following an evaluation of various publicly available PPI databases, we generated an integrated network merging high-performance transcriptomic and PPI datasets. Finally, we applied the PPR algorithm on this integrated network, *personalized* by a set of stalk cells marker training genes, to identify new genes with angiogenic function. The biological relevance of these predicted genes was validated through Gene Ontology (GO) [9] enrichment analysis, which confirmed their association with key processes driving angiogenesis and the pathological vascular transition essential for tumor growth.

## Materials and methods

### Personalized PageRank Algorithm

Historically, a web-surfer paradigm has been used to describe the PR algorithm which is based on random walks on graphs: a surfer randomly follows the links between webpages with a probability *α*, but at some point it may instead teleport to any random page with a probability 1 - *α*. This random teleportation is necessary to prevent the surfer from being “stuck” on a node without outgoing links or from running forever in a loop, which would be unlikely for a real web surfer. Mathematically, the PR algorithm can be formulated as follows:

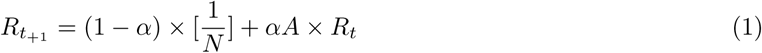

where *R_t_*_+1_ is the distribution of the importance scores at the time point *t* + 1, (1-*α*) is the probability to stop following the links and teleport with 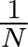 probability to any node, 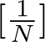 is the matrix of teleportation probabilities, *α* is the probability to follow the graph’s links, *A* is the network’s adjacency matrix and *R_t_* is the distribution of the importance scores at the time point *t*. This process repeats upon convergence of *R*. Another interpretation of the PageRank score *R* is the eigenvector of the graph’s adjacency matrix (rescaled so that the sum of each column equals 1) corresponding to the principal eigenvalue:

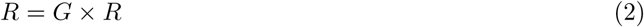

The PPR algorithm differs from the PR version in the way the distribution of the teleportation probability is formulated. Indeed, this distribution is no longer uniform as more weight is assigned to *prioritization nodes*, which allows the surfer to teleport to these nodes more frequently. From this perspective, the PPR algorithm is more relevant for the prediction of entities having properties similar to those of the prioritization nodes. In the context of gene function inference the PPR variant is more relevant than the original PR since we are interested in discovering new genes having a function similar to the prioritization nodes, thus the probability to teleport to these genes is higher than for the rest of the genes.

For the sake of clarity, the prioritization nodes will be referred to as “training genes”, which by definition are genes already known to be involved with a disease or biological process of interest serving as reference genes for the gene prioritization workflow.

### Training genes

50 genes having angiogenic stalk cell properties (S1 Table) previously curated [10] through single-cell RNA sequencing of 56,771 ECs from human, mouse and culture lung tumor models to investigate TEC phenotypes were used for inference as *prioritization nodes* for the PPR algorithm. However, due to the multiple data sources considered in this work, some training genes were simply absent from certain datasets, resulting in variations regarding the number of training genes actually present in the networks generated.

### Datasets

The networks for the PPR algorithm were constructed from publicly-available RNA-seq datasets and publicly available PPI databases. To infer cell-type-specific functions, specifically angiogenic stalk cell properties, we integrated transcriptomic data from EC subjected to angiogenic-relevant treatments. This integration was essential because standard PPI networks, while providing high-quality physical interaction data, are largely context-independent and lack the specificity necessary to distinguish cell type-specific regulatory programs.

### Protein-protein interaction databases

PPI databases, described in Table 1, were collected from publicly available resources, except for GeneCards [11] which was only available upon academic request (data from 2023). The PPI datasets were preprocessed differently because each database used its own *confidence score* (CS), necessitating normalization of the interaction weights in each one of them to ensure fair comparisons. In the BioGrid [12, 13] database (data from 2024), the ranges of these scores depended on the *Experimental System* value considered, which was a metric specific to this database. As a result, the scores were normalized per *Experimental System* value as follows:

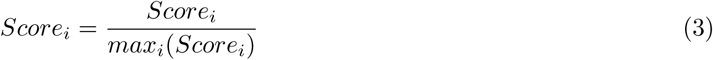

**Table 1.** PPI databases considered for building an integrated network.

| Database | Description | N nodes | N edges |
| --- | --- | --- | --- |
| GeneCards | Integrative database providing comprehensive, user-friendly information on all annotated and predicted human genes | 19,190 | 2,629,340 |
| STRING | Database of known and predicted protein-protein interactions | 19 611 | 3 726 381 |
| ConsensusPath | Collection of interaction networks from Homo Sapiens including binary and complex protein-protein, genetic, metabolic, signaling, gene regulatory and drug-target interactions | 18,412 | 5,240,113 |
| BioGrid | Biomedical interaction repository with data compiled through comprehensive curation efforts | 16,312 | 296,338 |

For the STRING [14–16] database (data from 2023), the CS values were ranging from 150 to 999. In order to ensure keeping relevant interactions from this database, we conducted a benchmark on filter values for these scores (starting from 100 to keep all the data and going up to 700 to filter as much data as possible while preserving a reasonable number of interactions). After determining the optimum threshold for the CS, they were normalized as described in Eq 3. In the ConsensusPath [17, 18] database (data from 2023), in addition to the *interaction confidence scores* we have also used the *number of publications* values as follows: in case the interaction confidence was not listed for a particular interaction, we have imputed it according to the average value of the score corresponding to the number of publications supporting the interaction. For the GeneCards database, the CS values were ranging from 1 to 9 and were normalized according to Eq 3.

### Omics datasets

Omics datasets, described in Table 2, which were used to create graphs based on correlation, were downloaded from Gene Expression Omnibus (GEO) [19, 20] and BioStudies [21]. From these bulk datasets, selected for comprising ECs subjected to angiogenic-relevant treatments, networks were created as follows. First, only datasets comprising more than eight samples were retained. Then, the poorly expressed genes and genes having a low variation were filtered out (using the 5^th^ percentile as a threshold), and finally the gene-gene correlation matrices were computed with the Pearson’s *r* correlation. Furthermore, as part of a hyperparameter fine-tuning step, different correlation thresholds (0.6, 0.7, 0.8, 0.9) were applied to the networks previously generated and analyzed with the PPR algorithm to find the best correlation threshold value.

**Table 2.** EC stalk cell-specific omics datasets considered for building an integrated network.

| Dataset | Source | Description | N samples | N genes |
| --- | --- | --- | --- | --- |
| E-MTAB-7774 | BioStudies | Deubiquitinase USP10 regulates Notch signaling in the endothelium | 12 | 23,549 |
| GSE45750 | Gene Expression Omnibus | Genome-wide analysis of differential gene expression of PFKFB3 under or overexpressed upon inhibition or activation of notch signaling in endothelial cells | 24 | 18,928 |
| GSE89174 | Gene Expression Omnibus | Genome-wide analysis of differential gene expression of contact-inhibited versus proliferating endothelial cells | 24 | 20,413 |
| GSE221860 | Gene Expression Omnibus | RNA-seq provides insights into VEGF-induced signaling in Human Retinal Microvascular Endothelial Cells | 12 | 22,553 |
| GSE77597 | Gene Expression Omnibus | RNA-sequencing of HU-VECs treated with Tie2 activating antibody | 8 | 24,846 |
| E-GEOD-18913 | BioStudies | Transcription profiling of HUVECs for Egr-3 knockdown in VEGF-treated cells | 24 | 13,051 |
| E-GEOD-37741 | BioStudies | Effects of Jmjd6 knock-down on human umbilical vein endothelials | 12 | 13,216 |

### Benchmark protocol

To evaluate the predictive performance of our approach, we conducted a comparative benchmarking against 1) established gene prioritization algorithms, such as SVM and RF, 2) renowned web-based gene prioritization tools such as *GeneMANIA* [22, 23] and the *ToppGene Suite* [24–26], 3) the original PageRank algorithm.

### Benchmarking against other gene prioritization algorithms

To ensure a rigorous and unbiased comparison, we computed a 5-fold cross-validation (CV) strategy using the same folds. Consistent with the optimization of the PPR model, we performed systematic hyperparameter tuning for both SVM and RF using the integrated network as input. This approach ensured that the comparison was based on the optimal configuration for each of the three gene prioritization methods, providing a fair assessment of their respective capabilities.

Both RF and SVM methods first required the conversion of the integrated network, initially in a graph format generated with the Python package *Networkx* [27], to an adjacency matrix representing all the connections between the genes from the network using the interaction weights as value. Then, labels were set based on whether the genes were among the training genes (label set to 1) or not (label set to 0). Therefore, the interaction weights were used as features and the labels as targets. Then, the 50 training genes were split in 5 folds of equal size in order to perform a 5-fold CV (each time with the same random state value to ensure reproducibility), a method whose evaluation metrics, including accuracy, AUC-ROC (also called AUC) and precision-recall, have been taken into account in order to determine the best hyperparameters for each algorithm.

To optimize the RF model, we performed a systematic grid search across three key hyperparameters: *N estimators* with values ∈ [1, 100, …, 500], *Max features* (either “Log2” or “None”) and *Max depth* with values ∈ [1, 100, …, 500]. This resulted in 6×2×6 = 72 potential configurations for the algorithm. For each configuration, a 5-fold CV was computed as follows: for each run, a RF model was fit on the training set (representing 80% of the training genes) and evaluated on the test set (representing 20% of the training genes). Accuracy, AUC and precision-recall were then calculated from this evaluation. After running this 5-fold CV, the average value for each of these metrics was determined, resulting in a global set of metrics for each potential hyperparameter configuration. Then, based on the best AUC, the optimum configuration was determined. Finally, using the best hyperparameters, a final 5-fold CV procedure was conducted to evaluate the optimized RF model performance using the resulting average AUC.

For the SVM model, we optimized three hyperparameters: *C* with values ∈ [0.1, 1, 10, 100, 1000], *gamma* with values ∈ [1, 0.1, 0.01, 0.001, 0.0001] and lastly *kernel* (either “rbf”, “poly”, “linear” or “sigmoid”). While we initially intended to utilize the *GridSearchCV()* function from the *Scikit-learn* [28] Python package for an exhaustive optimization, the high computational cost led us to adopt a more efficient approach utilizing the *RandomizedSearchCV()* function. The data were partitioned as described previously, and a grid search equal to a third of all the potential parameters combinations was set for the *RandomizedSearchCV()* function, leading to the same fit and test protocol as mentioned previously. This allowed for the identification of optimal hyperparameters within this restricted yet representative search space. Then, similarly as done with the RF model, the best hyperparameters combination was used to perform a final 5-fold CV to evaluate the optimized SVM model performance.

### Benchmarking against web-based gene prioritization tools

Initially released in 2010, GeneMANIA (”MANIA” standing for “Multiple Association Network Integration Algorithm”) is a bioinformatics web tool [https://genemania.org] which aims as predicting the function of query genes by searching thousands of functional association networks (co-expression, pathways, protein interactions, …) to find relevant and related genes. Similarly to the PPR algorithm, this tool relies on a *guilt-by-association* principle to infer the function of lowly characterized query genes based on their interactions with their well-studies neighbors.

GeneMANIA allows users to upload their own network prior to initiating the gene prioritization workflow. That is why we had initially planned to do so in order to make a fair comparison between this method and ours. However, likely due to the size of our network, this has been impossible as the page would load endlessly. Instead, we simply entered our 50 training genes in the input section and generated a model with the default parameters, resulting in a network made from 538 datasets (20 co-expression studies, 4 co-localization studies, 23 genetic interactions references, 6 pathways databases, 432 physical interactions studies, 51 predicted interactions collections and 2 shared protein domains databases). Afterwards, we retrieved the prioritized genes and evaluated their biological relevance and their novelty to compare this method against ours. Similarly, the ToppGene Suite is a web-based [https://toppgene.cchmc.org] collection of bioinformatics tool for gene prioritization. It offers users three types of analysis: 1) Gene Enrichment Analysis (through a tool called *ToppFun*) which aims as identifying pathways, biological functions and disease associations over-represented in the input list of genes; 2) Gene Prioritization (with a tool called *ToppGene*) by ranking a list of candidate genes (called “test set”) according to their functional similarities with a set of training genes (called “train set”); 3) Network Analysis (with a dedicated software named ToppNet) to prioritize genes based on their topological importance within PPI networks. We decided to opt for the latter, using as train set our 50 training genes and as test set the remaining genes in our network. We then had to chose a prioritization method between K Step Markov, Hyperlink-Induced Topic Search with Priors, and PageRank with Priors (which is the definition of PPR). Therefore, to ensure making a fair comparison with our method, we selected the third option as prioritization method with the same optimum damping factor value previously determined for our integrated network. Then, we retrieved the prioritized genes before assessing their novelty and biological relevance, therefore providing a thorough benchmark between GeneMANIA, ToppNet and our PPR implementation.

### Benchmarking against the original PageRank algorithm

To demonstrate the added value of the personalized variant in the context of gene prioritization, we compared it to the original version of the PR algorithm. To do so, we applied the PR algorithm on our network, without any training genes biasing the random walks, which resulted in a ranking of all the genes in the network based on their importance. Then, using the same cut-off as for the PPR version (see the Results section), we retrieved prioritized genes, whose biological relevance and novelty were afterwards assessed to compare this version of the PR algorithm against the other methods previously described.

### Assessing the novelty of the predicted genes via text mining

To assess the novelty of our findings, we developed a custom *text mining* algorithm, leveraging the *BioPython* [29] library, to perform automated, exhaustive queries of the Pubmed database [https://pubmed.ncbi.nih.gov/] in order to investigate the existing research landscape for potential predicted genes. The algorithm extracts the exact number of papers related to each query gene by reading the titles and abstracts available in the Pubmed database. By supplying specific keywords as input, the algorithm distinguishes between general biological knowledge and context-specific data. If no specific context is provided through a keywords prompt, the algorithm will look for general papers only. To ensure this PubMed search (PS) is exhaustive, all relevant aliases (extracted from GeneCards) of the input genes are considered. The resulting output table contains the number of papers (general and context-specific) and the related PubMed Identifiers (PMIDs) for each query gene, providing an overview of the current scientific knowledge about the input genes.

Using this number of papers, and thresholds determined internally, we identified two types of genes: 1) lowly characterized genes, having up to 50 general papers and 2) partially characterized genes, having up to 10 publications in a specific context yet more than 50 general papers (see the Results section). Interestingly, a gene well studied (i.e. with more than 50 general publications) can be poorly investigated in a specific context (such as angiogenesis in our case), making it an interesting candidate for further investigation in this topic.

### GWAS/TWAS cross-referencing

As an orthogonal, hypothesis-independent step, the top genes prioritized by the PPR algorithm were cross-referenced against publicly available human genetics resources. Genome-wide association evidence was retrieved from the NHGRI-EBI Genome-Wide Association Study (GWAS) Catalog [30]. Genes without a reported genome-wide significant association were further queried against transcriptome-wide association evidence using Transcriptome-Wide Association Study (TWAS) Atlas 2.0 [31], which links GWAS loci to candidate genes through genetically regulated gene expression. This allowed us to distinguish genes with prior support from human genetics studies from candidates entirely novel to the BfBio predictions.

### Assessing the druggability of the predicted genes

In novel target discovery, a major challenge is assessing its druggability which involves analyzing its structure, identifying potential binding sites for drug-like molecules and most of all predicting the likelihood of drug interactions. Nowadays, several computational tools have been created for this purpose.

### AlphaFold 3D structure predictions

Developed by Google DeepMind, AlphaFold [32, 33] (AF) is an AI program designed to predict the 3D structures of proteins based on their amino acid sequences using advanced deep learning techniques. It demonstrated a high accuracy in its predictions, notably through several predictions challenges over the last years, leading to a diversification of its applications over time, such as on a whole proteome [34] for instance. Even though AF3 [35], its latest update, is still in development and limited to academic and commercial uses only, AF 2 has been widely distributed as a web-server [https://alphafold.com], which provides the 3D structure (resolved or predicted) of any query protein or gene. Therefore, we greatly relied on this database to investigate the potential structure of our predicted genes since they were novel, meaning they had no officially resolved 3D structures yet.

### Protter

Until recently [36, 37], transmembrane proteins were considered more challenging than regular proteins in the context of drug design due to their flexibility, their tendency to aggregate and their instability [38]. That’s why inquiring about the potential transmembrane status of a target protein is a crucial step in drug design. The leading authority in this field is Protter [39], an open-source web tool for the visualization of phosphorylation site, isoform sequence variant and peptide identification to name a few. Taking in input either a UniProt protein accession number or an amino acids sequence, it then displays a sketch showing proteomics information such as variants, signal peptides, post-translational modifications (PTMs) and above all if the query protein effectively crosses the membrane. This web-tool is a valuable resource for conducting a preliminary druggability benchmark on a set of predicted potential novel targets.

### PockDrug

A major step in drug discovery, notably in the target identification phase, is the prediction of a protein pocket’s ability to bind ligands with a high affinity, which defines itself the druggability of the potential novel target considered. To this end, a web-server tool [https://pockdrug.rpbs.univ-paris-diderot.fr] named *PockDrug* [40, 41] has been created. It implements a well-established scoring function called *fpocket* [42, 43] While its main purpose is to predict pocket druggability, this can be achieved through two types of query. In the first case, the input protein already has an estimated interaction pocket, whose druggability probability will then be evaluated by PockDrug. This is a reliable mean to confirm (or disconfirm) a previous pocket prediction. In the second case, should the only information available be the protein’s structure, the druggability prediction can also be estimated. Due to the novelty of our predicted targets and their limited associated information, we relied on this second option to assess their druggability probability. Both methods produce an output table listing relevant information about the predicted pocket, such as its volume, surface or number hydrophobic residues to name a few. All these metrics are then aggregated into a druggability score (ranging from 0 to 1), evaluating the therapeutic potential of the target considered.

### PROST

A major obstacle in drug design is the lack of resolved 3D structure for a potential target, as it makes it impossible to accurately model the pocket, which as a result hinders drastically the understanding of the ligand/protein interaction. A workaround to deal with this issue is *homology modeling* [44–46], a computational method to predict the 3D structure of a lowly characterized protein based on known structures of similar proteins which then acts as templates. In this way, potential binding sites for drug-like molecules, interactions and protein functions can be unraveled.

In recent years, a number of homology modeling tools have emerged, such as *SWISS-MODEL* [47], *I-TASSER* [48] and *PRIMO* [49]. In 2022 a new homology method was created: *Protein Language Search Tool (PROST)* [50], a web-server for homology prediction tasks incorporating a language model and a quantization technique to preserve evolutionary information when representing proteins in a numerical format. Even though AF2 already provides an homology tool for finding similar proteins, PROST can be used as a complementary option when no evident homology is available for a target gene.

## Results

### Building an integrated EC stalk cell-specific interaction network

After collecting several omics and PPI datasets, we conducted a benchmark to assess their individual performance in order to integrate the best PPI database to the best omics datasets, therefore resulting in a reliable and EC stalk cell-specific biological network. To do so, the PPR algorithm was first applied to each dataset separately. To maximize the predictive accuracy of the transcriptomic networks, we systematically explored the parameter space by optimizing the *correlation threshold r* and the *damping factor α*. The AUC performances were computed by running the PPR algorithm with a 5-fold CV for different damping factors *α* (ranging from 0.1 to 0.9). The results were then averaged across the 5 folds and are depicted in Table 3.

**Table 3.** Omics datasets performance with their best hyperparameters, including Pearson’s correlation *r* and damping factor ***α***.

| Dataset | N nodes | N edges | N training genes | AUC | $r$ | $\alpha$ |
| --- | --- | --- | --- | --- | --- | --- |
| E-MTAB-7774 | 23,549 | 19,036,244 | 44 | 0.78 | 0.6 | 0.1 |
| GSE45750 | 20,988 | 2,990,101 | 45 | 0.75 | 0.7 | 0.8 |
| GSE89174 | 21,769 | 7,385,961 | 48 | 0.70 | 0.7 | 0.7 |
| GSE221860 | 22,553 | 5,386,491 | 47 | 0.77 | 0.8 | 0.6 |
| GSE77597 | 24,846 | 20,482,661 | 47 | 0.71 | 0.7 | 0.9 |
| E-GEOD-18913 | 13,051 | 7,560,761 | 42 | 0.68 | 0.6 | 0.1 |
| E-GEOD-37741 | 13,216 | 2,171,258 | 38 | 0.62 | 0.9 | 0.7 |

Fig 1 summarizes the impact of the damping factor *α* and the correlation threshold *r* on the AUC for omics datasets, whereas Fig 2 shows the PPR performance for the PPI databases according to several damping factor values. Interestingly, one can notice that for all the PPI databases, the PPR algorithm performance dropped with the increase of the damping factor. For the STRING PPI database, we were looking for the best combination of the confidence score threshold (ranging from 100 to 700) and the damping factor unlike the other PPI databases where all interactions were used. Following this optimization step, we merged the highest-performing omics and PPI networks into a single *integrated network*. We then applied the PPR algorithm on this network, using the 50 angiogenic stalk cells training genes, to drive the personalization and generate final predictions.

**Fig 1.**
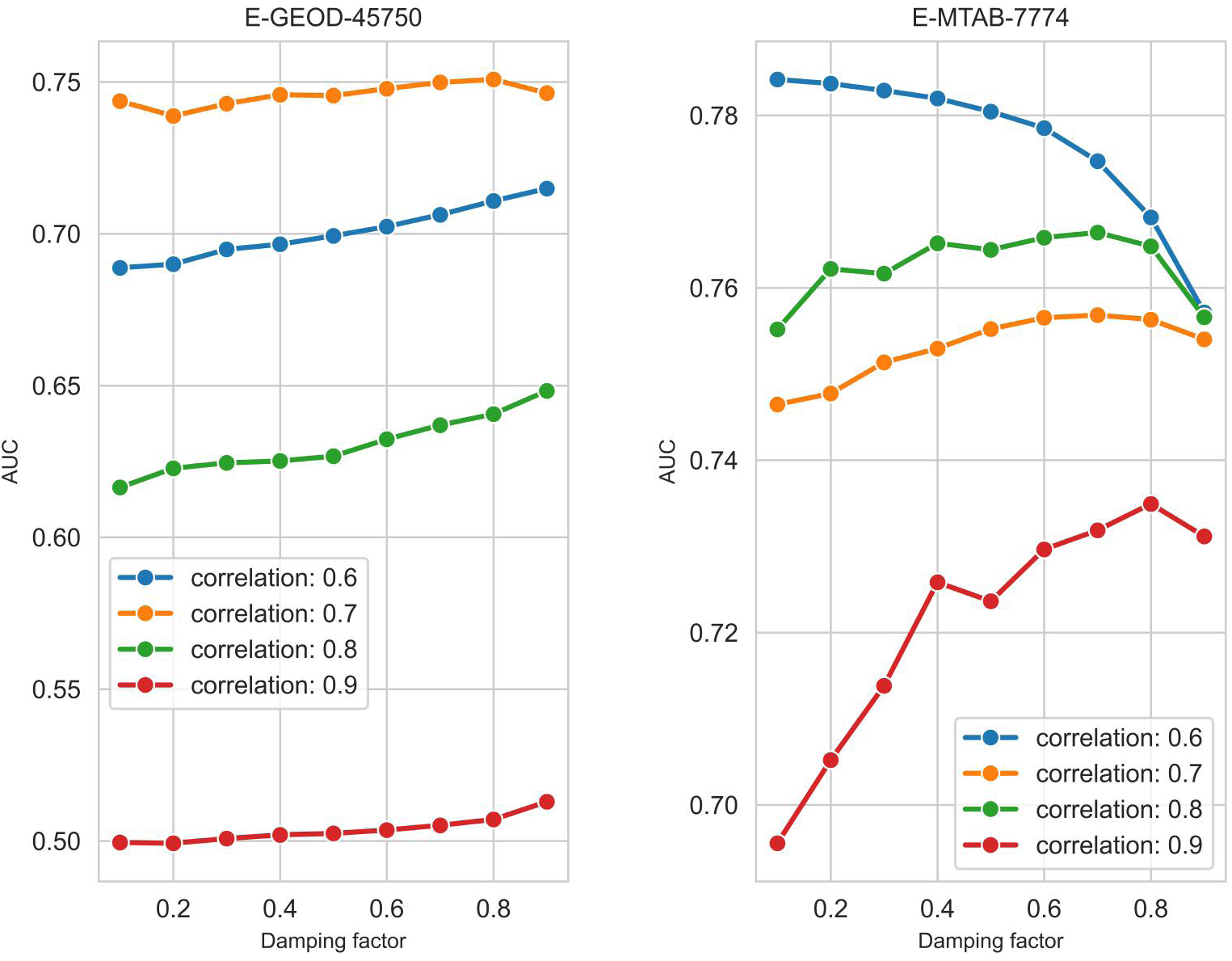
Omics datasets E-GEOD-45750 (*left*) and E-MTAB-7774 (*right*) performance across different correlation thresholds and values of damping factor ***α***

**Fig 2.**
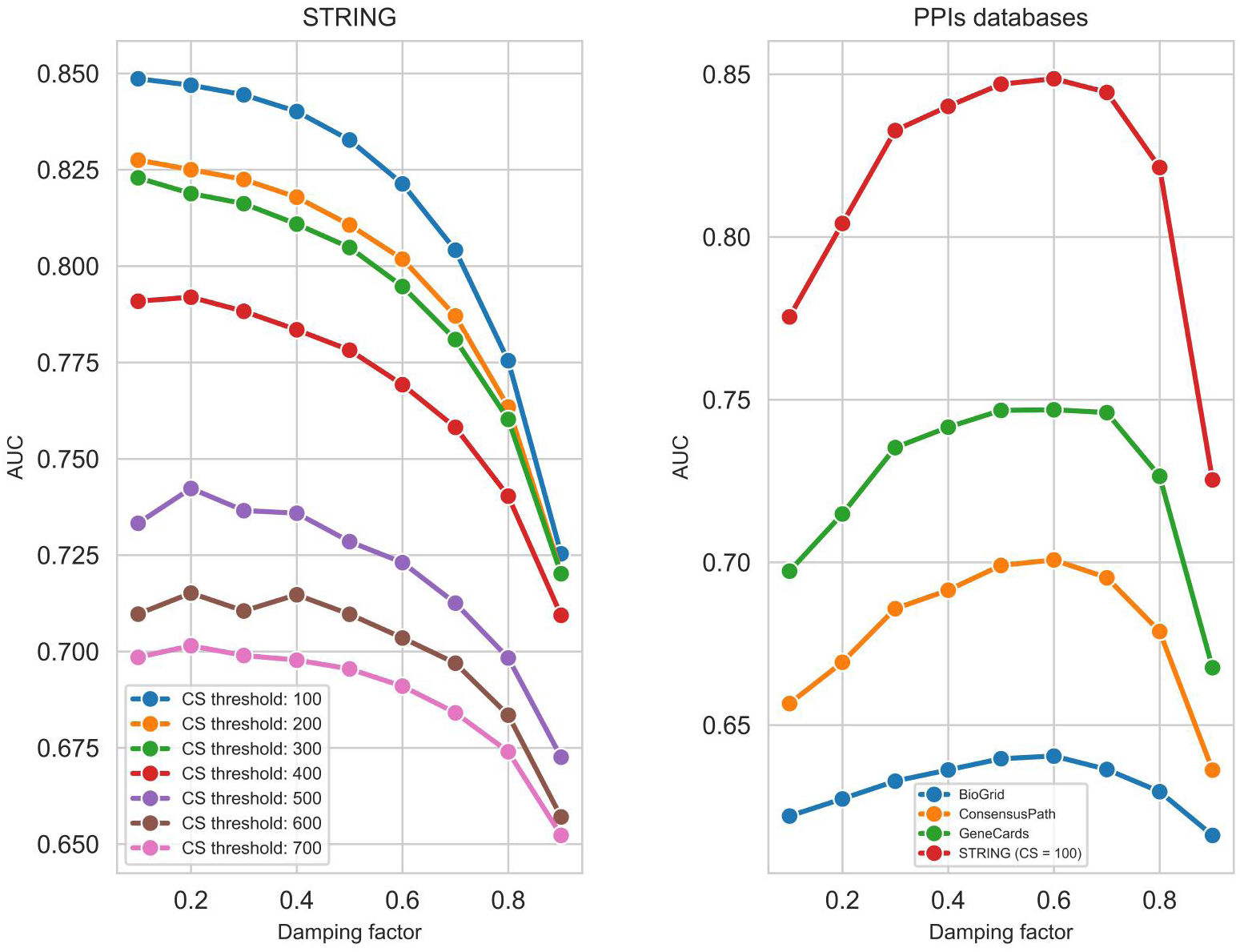
PPI databases performance across different values of damping factor ***α***

For the PPI databases, the best results were obtained with STRING and the following hyperparameters: *α* = 0.1 and *CS* threshold = 100, as shown in Table 4.

**Table 4.** PPI databases performance with their best hyperparameters, including confidence score (*CS*) threshold (for STRING only) and damping factor ***α***.

| Dataset | N nodes | N edges | N training genes | AUC | $CS$ threshold | $\alpha$ |
| --- | --- | --- | --- | --- | --- | --- |
| STRING | 19,622 | 6,587,702 | 49 | 0.85 | 100 | 0.1 |
| GeneCards | 19,190 | 2,629,340 | 49 | 0.75 | / | 0.2 |
| ConsensusPath | 18,412 | 5,240,113 | 48 | 0.70 | / | 0.6 |
| BioGrid | 16,360 | 296,676 | 43 | 0.64 | / | 0.6 |

For the omics datasets, the best performance was achieved with the dataset E-MTAB-7774 with *r* = 0.6 and *α* = 0.1, followed closely by the dataset GSE45750 with *r* = 0.7 and *α* = 0.8.

From all datasets (both omics and PPI), the STRING database showed the best AUC score. Thus, we computed two *integrated networks* made from the combination of STRING (with a *confidence score* threshold = 100 and *α* = 0.1) and the 2 best omics datasets, E-MTAB-7774 (with *r* = 0.6 and *α* = 0.1) and GSE45750 (with *r* = 0.7 and *α* = 0.8). The integrated graph made from the combination of STRING and E-GEOD-45750 showed better performance than the individual datasets, comforting us about the need to add EC-specific information to the STRING PPI database (Table 5; Fig 3).

**Fig 3.**
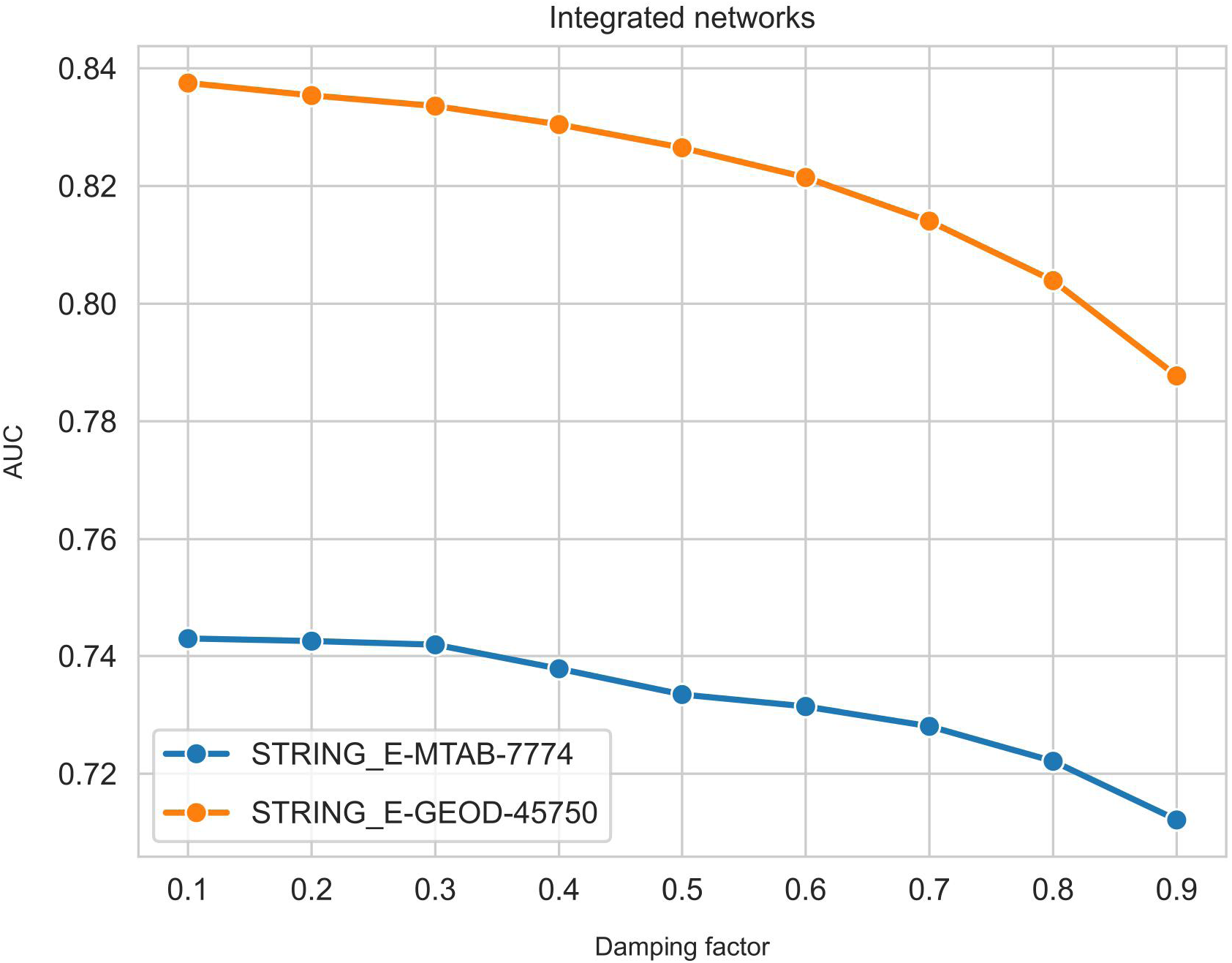
Performance of the integrated networks across different values of damping factor *α*

**Table 5.** Best PPR performances for the two integrated networks according to the AUC.

| Integrated network | $\alpha$ | AUC | N genes | N edges | N training genes |
| --- | --- | --- | --- | --- | --- |
| STRING + GSE45750 | 0.1 | 0.837 | 23,680 | 9,675,504 | 49 |
| GeneCards + E-MTAB-7774 | 0.1 | 0.742 | 26,449 | 19,243,243 | 49 |

The damping factor $\alpha$ has been re-optimized after integrating each dataset.

### Predictions

Given the robust performance of the integrated graph, we proceeded to predict angiogenic stalk cell genes using the 50 genes described in previously as personalization vector for the PPR algorithm. A significant challenge in this approach is that the PPR output is a continuous ranking of all the genes in the input network, from high to low *PageRank score*, corresponding to a distribution summing up to 1. This ranking does not inherently provide a defined threshold for identifying the most biologically relevant candidates. To address this problem, we performed a set of permutations and compared the distribution of the *PageRank score* from the real data with the distribution of the *PageRank score* from the permuted dataset. These permutations were computed in three different ways 10 times each:

1. A graph with randomly-generated edges and 50 randomly selected from the whole network (except the actual 50 previously described genes) training genes used as personalization vector for the PPR algorithm.
2. A graph with randomly-generated edges with the 50 real training genes used as personalization vector for the PPR algorithm.
3. The actual integrated network previously described with 50 randomly selected training genes used as personalization vector for the PPR algorithm.

To ensure preserving the same number of nodes, edges, and the overall degree distribution, the graph shuffles described in 1) and 2) were performed by simply randomly swapping the genes names, resulting in randomly-generated networks with the same structure as the original integrated network.

By visualizing these distributions in a *log*_10_ scale (Fig 4), we identified a threshold designed to minimize the inclusion of random *PageRank score* distributions while maximizing the retention of high-scoring genes in the actual *PageRank score* distributions. This analysis led us to the selection of a threshold *log*_10_ = −4.4, corresponding to a *PageRank score* ≈ 4.10*^−^*^5^. This cutoff prioritized the top 49 ranked genes (S2 Table) within the integrated network for further analysis.

**Fig 4.**
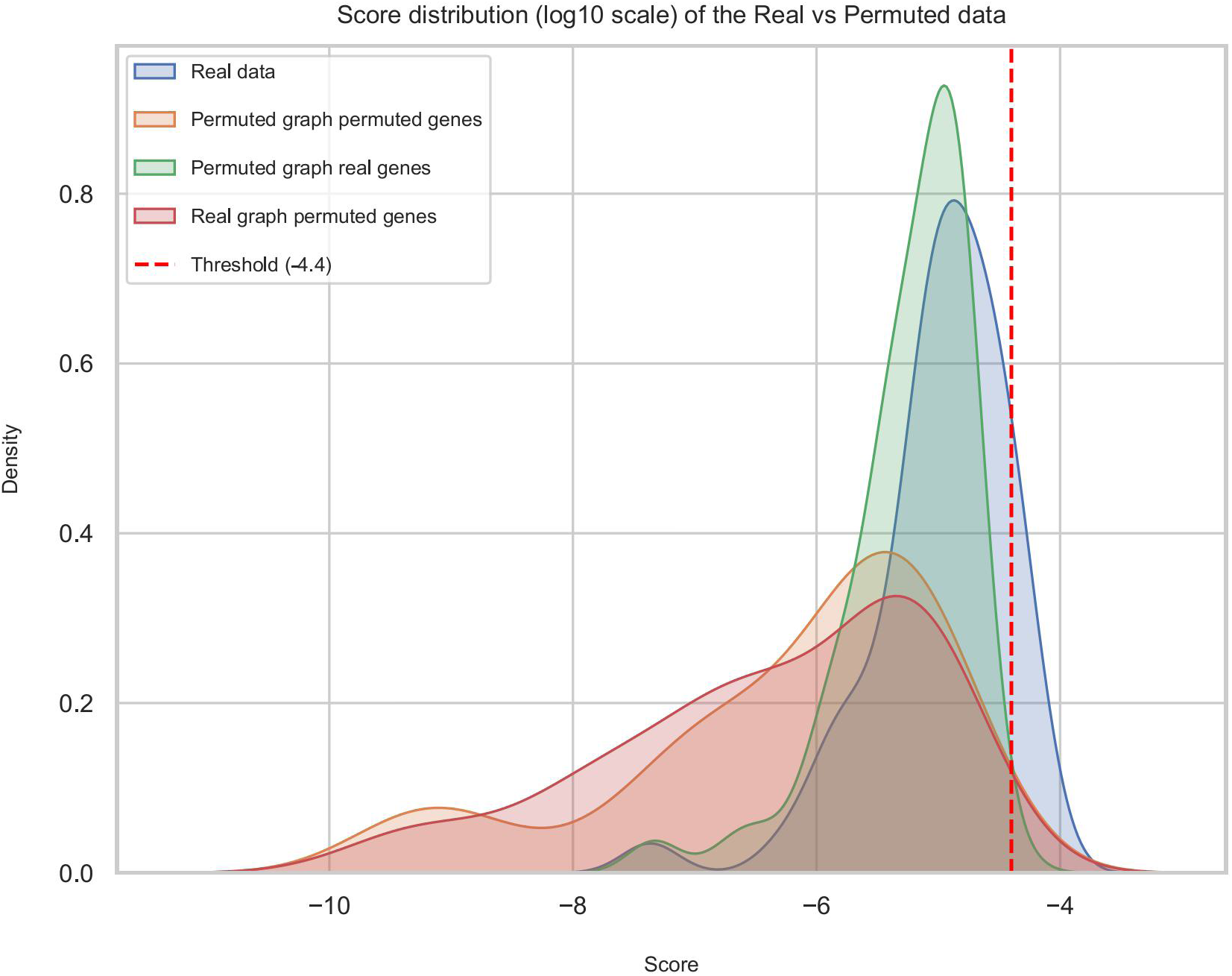
Distribution of PageRank scores from the real data vs from different types of permuted data

### Gene Set Enrichment Analysis

To investigate the potential biological relevance of these 49 predicted genes, we performed a Gene Set Enrichment Analysis (GSEA) with Enrichr [51–53], a GSEA web-based tool that contains hundreds of thousand annotated gene sets [https://maayanlab.cloud/Enrichr/]. The purpose of this method is to assess whether genes from a specified set are disproportionately involved in a phenotype or biological process of interest. The results showed that these predicted genes are significantly enriched in GO terms directly associated with angiogenesis processes, such as “Positive Regulation of Cell Migration”, “Regulation of Cell Migration”, “Positive Regulation of Cell Motility”, “Regulation of Endothelial Cell Proliferation” and “Regulation of Cell Population Proliferation” (Fig 5).

**Fig 5.**
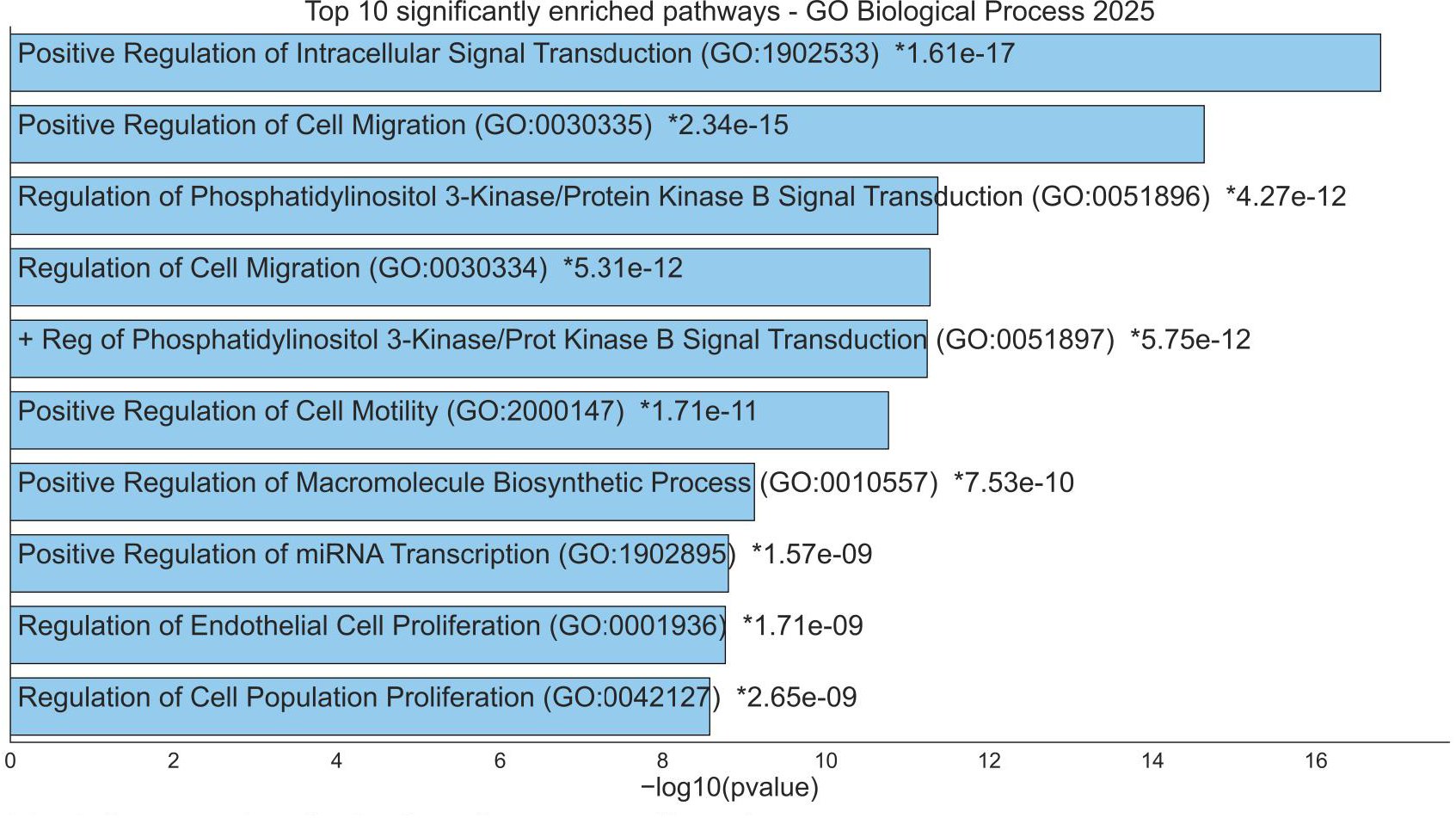
Enrichment Analysis for the 49 predicted genes.

Notably, we observed a substantial functional convergence between the training set and the predicted genes. Out of 45 GO terms significantly enriched in the training set (Fig 6), 23 were also significant for the top 49 predicted genes. This approximately 51% overlap in enriched biological pathways indicates that our network-based approach effectively expanded the stalk cell gene set while maintaining the core functional characteristics of the known regulatory landscape.

**Fig 6.**
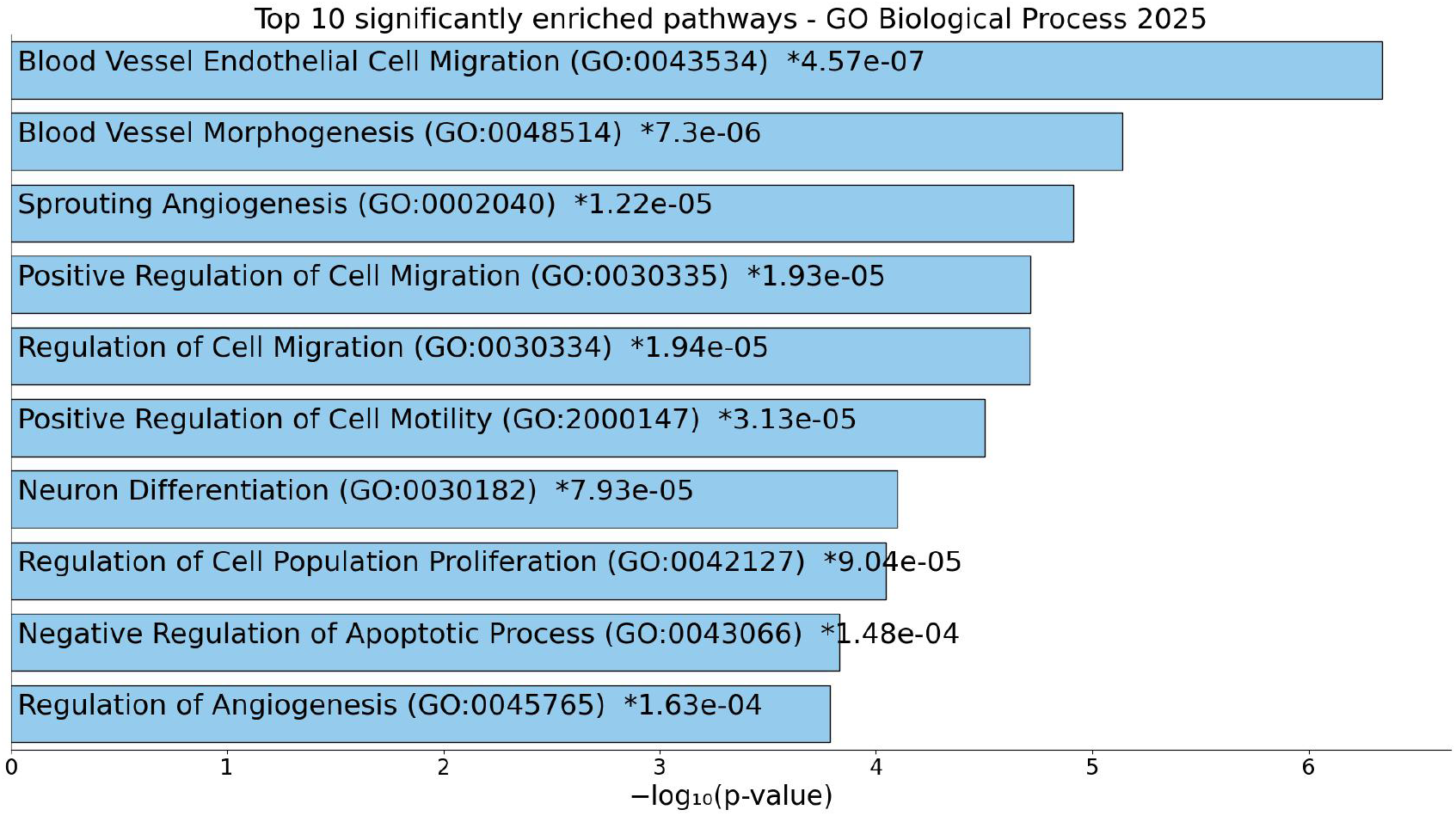
Enrichment Analysis for the 50 training genes.

Furthermore, despite identifying 4 lowly characterized stalk-like genes (see below), genes already largely associated with angiogenesis, and more particularly with the differentiation of EC into stalk cells, have also been identified by our algorithm. Indeed, among these top 49 predicted genes are 3 genes that have been described in the literature as contributing to this stalk cell phenotype: *ACVRL1* [54], *TIE1* [55] and *KDR* [56]. Therefore, these findings validate our model and further confirm the biological relevance of our computational predictions.

### Comparing our method with other gene prioritization algorithms

Once the 3 models (PPR, RF and SVM) were optimized (Table 6), we compared their predictive performance through a 5-fold CV process. To ensure a rigorous and unbiased comparison, we used identical 5 folds with fixed *random state* value set at the time of the splitting with the *KFold()* function from the *Scikit-learn* package. We selected the AUC as the primary evaluation metric to assess the prediction performance of each model (Table 7).

**Table 6.** Optimized hyperparameters for RF and SVM models.

| Method | Parameter 1 | Parameter 2 | Parameter 3 |
| --- | --- | --- | --- |
| RF | $N_{estimators} = 400$ | $Max\_features = \log 2$ | $Max\_depth = 100$ |
| SVM | $C = 0.1$ | $gamma = 0.1$ | $kernel = rbf$ |

**Table 7.** PPR, RF and SVM performance with a 5-fold CV.

| Method | AUC 5 fold-CV |
| --- | --- |
| PPR | $0.837 \pm 0.044$ |
| RF | $0.807 \pm 0.024$ |
| SVM | $0.558 \pm 0.047$ |

The benchmarking results demonstrate that our PPR algorithm outperforms RF and SVM in prioritizing angiogenic stalk cell relevant genes within our integrated network.

### Comparing our method with established web-based gene prioritization tools

After running a gene prioritization workflow with GeneMANIA using our 50 training genes as input data, which led to the generation of an integrated network made from 538 data sources, 20 genes were prioritized (Table 8).

**Table 8.** 20 genes prioritized by GeneMANIA using our 50 training genes as input.

| Gene | Score |
| --- | --- |
| ELK4 | $1.48650 \times 10^{-2}$ |
| VWF | $1.40761 \times 10^{-2}$ |
| CDH5 | $1.35098 \times 10^{-2}$ |
| GJA4 | $1.29727 \times 10^{-2}$ |
| TIE1 | $1.22562 \times 10^{-2}$ |
| PDGFRB | $1.17660 \times 10^{-2}$ |
| LAMA5 | $1.15151 \times 10^{-2}$ |
| IGFBP5 | $1.15067 \times 10^{-2}$ |
| DDIT4L | $1.11256 \times 10^{-2}$ |
| TSC22D3 | $1.10132 \times 10^{-2}$ |
| RAMP2 | $1.09295 \times 10^{-2}$ |
| ADGRL4 | $1.08792 \times 10^{-2}$ |
| IGFBP7 | $1.08322 \times 10^{-2}$ |
| KDR | $1.08310 \times 10^{-2}$ |
| TIMP3 | $1.08115 \times 10^{-2}$ |
| ESAM | $1.04502 \times 10^{-2}$ |
| PTMS | $1.03753 \times 10^{-2}$ |
| LDB2 | $1.03073 \times 10^{-2}$ |
| SP1 | $1.00855 \times 10^{-2}$ |
| RPSA | $1.00405 \times 10^{-2}$ |

Afterwards we performed a GSEA with Enrichr to investigate the biological relevance of these prioritized genes and verified if we could observe a convergence between the training genes and this set of genes. Out of 45 GO terms significantly enriched in the training genes, 13 were also significant for these 20 predicted genes (Fig S1), resulting in an overlap of approximately 31%, which is less than the 51% overlap observed with the 49 genes predicted with our PPR method. Furthermore, using our PS algorithm with “endothelial”, “angiogenesis” and “cancer” as keywords, only one gene with up to 50 general publications (therefore lowly characterized) was identified, compared to 5 with our PPR algorithm (see below). These results demonstrate better gene prioritization performances from our method.

Regarding ToppNet, after computing a PPR gene prioritization as explained previously, we used the same threshold we have determined with our permutations process to retrieve the first 49 genes ranked by ToppNet (S3 Table), and we assessed their biological relevance through another GSEA. We observed this time an even lower overlap between these 49 predicted genes and the training genes, as only 10 out of the 45 relevant GO terms significantly enriched in the training genes were found in the predicted genes (approximately 24%; Fig S2). However, a PS inquiry with the same keywords and filtering criterion revealed 3 potential understudied genes. Interestingly, it appeared 5 genes were commonly prioritized between ToppGene and our PPR algorithm: *EGFR*, *KRAS*, *MYC*, *TP53* and *FN1*. These results also demonstrate significant better gene prioritization performances from our method, even though they are still an improvement over those obtained with GeneMANIA regarding the identification of potential unexplored genes.

### Comparing our method with the original PageRank algorithm

To ensure a thorough comparison between these network-based gene prioritization methods, we applied the original version of the PR algorithm on the integrated network we generated earlier. In a similar way as ToppNet and the PPR variant, the original PR ranks all the genes in the network according to their importance and connectivity. Thus, we relied on the same threshold, the top 49 genes, to make further analyses. These 49 PR-ranked genes (S4 Table) were used to perform a GSEA to investigate their biological relevance. Among the 45 relevant GO terms significantly enriched in the 50 training genes, 11 were also enriched in these predicted genes (Fig S3), resulting in an overlap of approximately 31%. Furthermore, after running the same PS as detailed previously, 2 potential lowly characterized genes were identified. Table 9 summarizes the benchmark results between the 4 network-based gene prioritization methods. Overall, our PPR algorithm demonstrates better performances in the identification of biological relevant yet poorly investigated genes over established gene prioritization tools and algorithms, granting us confidence in the genes we have predicted.

**Table 9.**
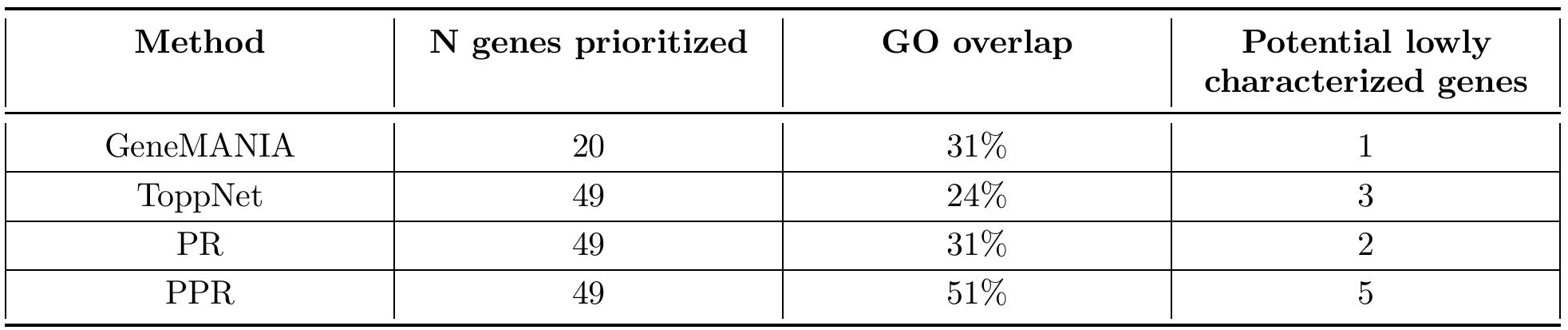
Benchmark results between GeneMANIA, ToppNet, PR and PPR.

### Characterization of lowly annotated candidate genes

Applying the PS algorithm to the 49 predicted genes with our PPR method, using “endothelial” and “angiogenesis” as keywords, we identified five lowly characterized genes. After excluding one non-coding RNA gene, we focused on the four remaining candidates: *PLPP4*, *ANKRD36B*, *FAM124A* and *FHIP2A* (also known as *FAM160B1)*, respectively ranked 1^st^, 3^rd^, 42^nd^ and 44^th^ out of the top 49 genes in the whole integrated network comprising 23,680 genes. S5 Table describes the PS results for the 49 predicted genes. To investigate the disease relevance and therapeutic potential of these four candidates, we leveraged the transcriptomic data on Disign Atlas [57]. All four genes were up-regulated across several cancer-related diseases, suggesting a potential involvement in oncology (Fig 7 showing the top 10 studies with the highest Log2 fold change values). Interestingly, except for *FAM124A*, these potential target genes are all up-regulated in Central Nervous System (CNS) phenotypes. *PLPP4*, *ANKRD36B* and *FHIP2A* additionally showed CNS-related disease signatures in this analysis (S6 Table); *FAM124A*’s DiSignAtlas profile was instead cancer-and metabolism-associated though its GWAS associations point to a CNS link via in independent resource.

**Fig 7.**
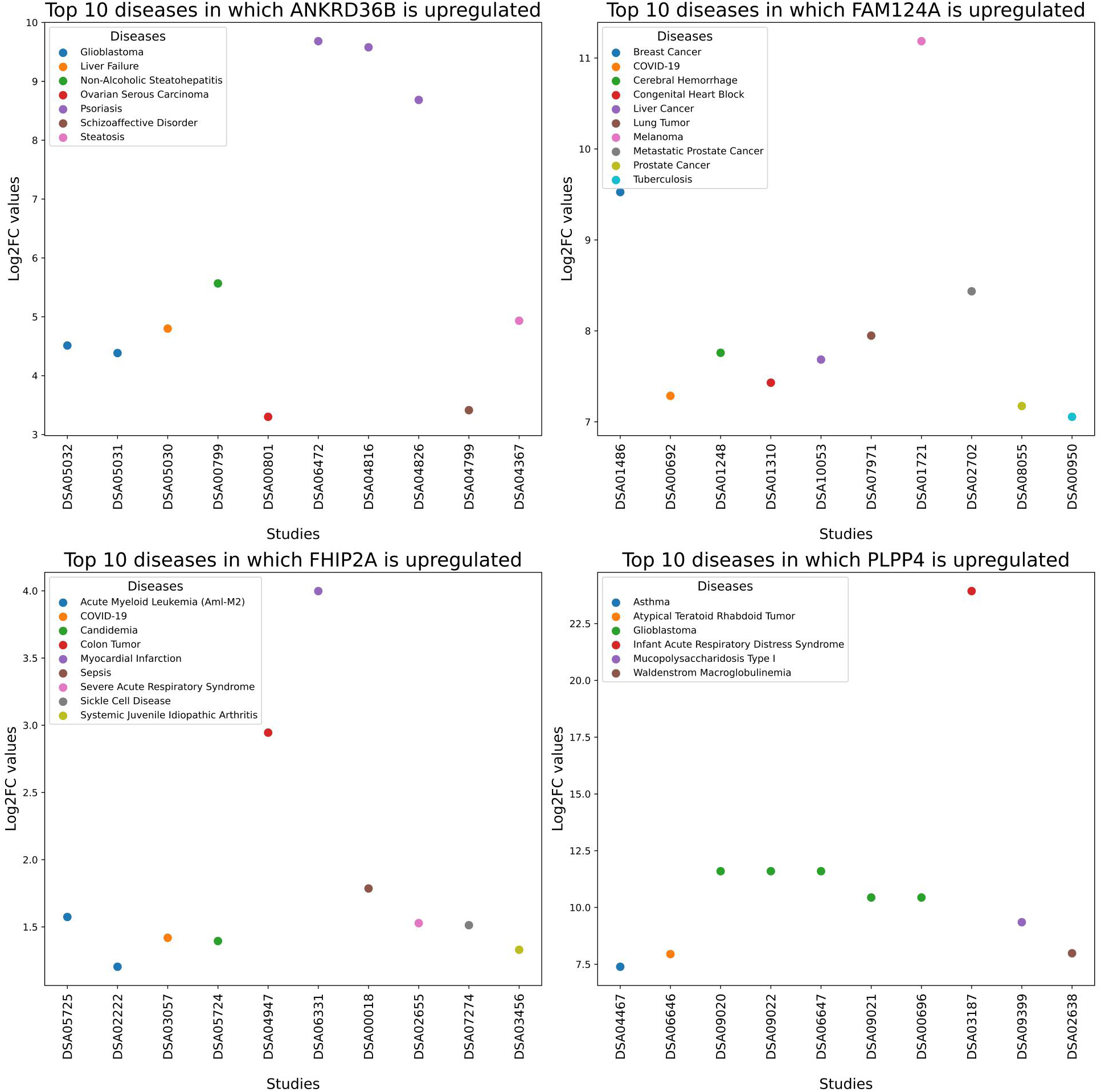
Top 10 diseases in which the four target genes are up-regulated.

As an orthogonal, hypothesis-independent line of evidence, we further examined whether these four candidates had been implicated in genome-wide or transcriptome-wide association studies. *PLPP4*, *FHIP2A* and *FAM124A* each carried GWAS associations converging on neurological and cognitive phenotypes [58–62], emphasizing the CNS signal seen for these genes in DiSignAtlas. *ANKRD36B*’s profile diverged from the other three: its largest cluster of GWAS associations related to socioeconomic and occupational measures, alongside respiratory function and metabolic traits, and it carried an independent TWAS association with malignant pancreatic neoplasm. None of the associations recovered from either resource for any of the four candidates were directly related to angiogenesis or vascular traits.

Among these candidates, *PLPP4* had the clearest existing functional annotation: according to GeneCards, *PLPP4* plays a role in phospholipid signaling [63], a pathway previously linked to endothelial migration, proliferation, permeability, and angiogenesis [64], providing a plausible mechanistic bridge between this candidate and the phenotype BfBio was designed to predict, despite its otherwise limited characterization. In summary, this multi-resource profiling identifies four previously uncharacterized genes with plausible relevance to EC function: consistently up-regulated across cancer-related disease datasets, converging on CNS phenotypes in three of four cases via two independent resources, and largely unexplored either structurally or in the specific context of angiogenesis.

### Preliminary druggability results

Considering how understudied these 4 target genes are, it is not surprising that no 3D structures were available in UniProt [65] at the time of writing. Still, to further investigate their potential druggability, we collected for each gene its AF3 predicted 3D structure (Fig S4). Then, combining Protter, PockDrug and PROST, we have drawn up a preliminary assessment of these four genes, which will be a good starting point for further investigation in the hope of conducting a full drug design process should these genes be promising. Table 10 provides an overview of relevant features we could identify for these 4 undercharacterized genes.

**Table 10.** Summary of druggability-related features for the 4 potential lowly characterized genes prioritized by our PPR algorithm.

| Gene | Average pLDDT | Transmembrane | Druggability probability | Homology | p-value |
| --- | --- | --- | --- | --- | --- |
| PLPP4 | 91.MrsMr12 | Yes | 0.98 | P0C8X4 | 1.34E-07 |
| ANKRD36B | 90.44 | No | 0.11 | P42048 | 1.86E-07 |
| FAM124A | 80.00 | No | 0.8 | P85251 | 1.28E-06 |
| FHIP2A | 79.69 | No | 0.95 | P0DSE3 | 2.88E-07 |

The PROST homology prediction results were calculated for each protein on the best predicted pocket from PockDrug rather than on the whole predicted 3D structure from AF3 to ensure biological relevance of the resulting binding pose in later drug discovery stages. At the time of writing, after gathering information from GeneCards, UniProt and DrugBank [66], a comprehensive drug-target database, no information related to existing drugs (approved or experimental) associated with these genes and their homologs could be retrieved, highlighting the novelty of our findings.

Even though *PLPP4* displays the best average predicted Local Distance Difference Test (pLDDT) value, an evaluation metric from AF which evaluates its confidence in the predicted 3D structure, according to Protter it seems to display 6 transmembrane domains (Fig S5), which might complicate the upcoming drug design steps as it would require to mainly focus on the extracellular domains instead of the whole protein. As a result, it would be best to shift the focus on one of the 3 remaining proteins. Despite exhibiting the lowest average pLDDT (even though according to AF such a score can be linked to a high confidence in the predicted structure), *FHIP2A* is associated with the best druggability probability score (computed by PockDrug) and the best p-value related to its best available homology structure. Although these results are preliminary, they pave the way for further investigation (including functional validation then drug discovery), and are reliable filtering criteria to invest more time in *FHIP2A* than in the other genes.

## Discussion

### Identification of potential lowly characterized genes associated with an angiogenic stalk cell phenotype

In this study, we applied a PPR algorithm to an integrated network combining a stalk cell specific omics dataset with a protein-protein database to infer novel genes involved in angiogenic stalk cell phenotype as the identification of such genes has a significant therapeutic importance for angiogenic therapies. Our model was personalized using 50 genes, curated from a single-cell RNA sequencing of human and murine angiogenic stalk endothelial cells within the lung tumor microenvironment.

Our results demonstrate that the integration of co-expression from omics datasets and physical interactions from PPI databases has a synergistic effect, with the AUC of the integrated graph outperforming the AUC of the individual graphs.

To address the challenge of thresholding continuous distributions of *PageRank scores* and retrieve the most promising genes in the network, we compared the distribution of the real *PageRank scores* against those of permuted data. This framework identified an optimal threshold (*log*_10_ = −4.4), prioritizing the 49 top ranked genes in the network. Validation through a GSEA confirmed a high degree of functional coherence between the training genes representing known regulators and our novel predictions (leading to a 51% overlap).

Based on a thorough benchmark protocol, our method has outperformed five other established gene prioritization tools regarding predictive abilities, which led to higher AUC and enrichment overlap for our PPR algorithm.

Finally, through an automated text mining, we identified four poorly investigated genes, particularly in the context of stalk cell/angiogenesis. These genes represent prime candidates for further experimental characterization to elucidate their role in angiogenesis and the tumor microenvironment, which is already hinted by their up-regulation in several cancer-related diseases. Given their high predictive scores and lack of prior characterization, these identified genes might constitute a potential therapeutic reservoir for the development of first-in-class anti-angiogenic strategies in oncology, pending further functional validation.

### Linking the candidate genes to potential diseases

The GWAS/TWAS cross-referencing analysis provides an additional, orthogonal line of validation for the prioritized genes. The tau-pathology signal across *PLPP4* and *FHIP2A*, together with the CNS-tumor and memory-performance associations for *FAM124A*, is a striking convergence with the CNS disease pattern already observed via DiSignAtlas. *ANKRD36B*’s association profile to respiratory and metabolic traits, with an independent TWAS link to pancreatic cancer, suggests it may act through a different biological axis than the other three candidates, warranting separate follow-up rather than being grouped with the CNS-associated genes. As larger GWAS and molecular Quantitative Trait Locus resources become available, additional human genetic evidence may further clarify the relevance of these computationally prioritized candidates to disease. Integrating network propagation with human genetics, therefore, represents a promising strategy for prioritizing genes for experimental validation.

### Provision of a new text mining tool to access the PubMed database

In the course of this project, it has become clear that a more efficient process is needed for literature search, which was initially conducted manually. Although this would ensure the papers we selected were completely relevant, this task had proven to be time-consuming. Inspired by the *Biopython* package, which provides a direct connection to the PubMed servers, we identified the functions allowing this connection and tweaked them to build our own script in order to finally automate large PS. Remaining true to the spirit of *Biopython*, which aims to make bioinformatics accessible to Python programmers, we designed our algorithm to be interactive (through a question/input prompt system) and very explicit so people who are not familiar with Python programming can still use it easily. As a result, in addition to our gene prioritization method, we provide an open-source text mining tool to explore the PubMed database, which proved useful for our work and allowed us to identify poorly characterized genes with relevant biological properties among our prioritized genes.

### Limitations of the study

Nevertheless, in its current state, BfBio has a few limitations that users should bear in mind. Indeed, on the one hand, due to the context-independent nature of the PPI databases, incorporating relevant transcriptomics datasets is crucial to ensure constructing a meaningful network in order to perform a reliable gene prioritization. However, in our work some of these bulk datasets displayed relatively small samples sizes, which may not fully represent the co-expression interactions needed for the network. Noteworthy, these samples were largely *in vitro*, a biological condition in which ECs tend to lose their *in vitro* phenotypes. On the other hand, gene prioritization depends heavily on the amount of information available from the literature, which in the context of novel target discovery leads to a paradox: if there are too many publications related to a gene then it cannot be considered as poorly characterized, losing its novelty, but if on the contrary there are none then the gene, being absent from the PPI databases, will most likely not appear in the integrated network (unless it is present in the transcriptomics datasets, highlighting the importance of incorporating this co-expression data to the physical PPI data) and consequently might be missed by the PPR algorithm during the gene prioritization. Also, users should be aware that the PS tool is limited to titles and abstracts, meaning the number of publications returned by the algorithm is the outcome of a very limited search space and may not represent the full extent of the available literature for the query genes.

In parallel of identifying potential novel genes involved in an angiogenic stalk cell phenotype, we had initially hoped to infer this function to already known genes by finding, through our PS tool, prioritized known genes with a limited number of publications but not yet reported to be involved in stalk cell phenotypes. Unfortunately, according to our PS results the genes predicted were either completely uncharacterized or already characterized in angiogenic/stalk cell pathways.

## Conclusion

Here we introduce Brain-for-Biotech, a framework designed to identify genes important for vascular ECs, which are crucial cells for angiogenesis, vascular homeostasis, hemostasis and blood/tissue barrier function but also critical mediators of immunity and cancer progression. BfBio utilizes a PPR algorithm on an integrated network of different omics datasets and publicly available gene-gene/protein-protein interaction databases. In this study, we apply BfBio’s predictive capabilities to infer angiogenic stalk cell phenotype function in genes for which this function was not known before. By leveraging a set of genes characterizing the stalk cell cluster in lung tumor EC models previously identified, we have achieved a high AUC (0.837) performance and have even outperformed five other gene prioritization methods. Enrichment analysis, coupled with a text mining application, further confirmed that among the 49 predicted genes four of them were poorly characterized yet possessed biologically relevant properties and were linked to cancer, thereby validating BfBio as a robust tool for prioritizing novel therapeutic targets in vascular biology. In addition to providing a reliable open-source gene prioritization algorithm, we have also provided an powerful Python text mining tool to help researchers with their literature search with PubMed. Finally, although this study primarily focused on stalk cell genes and angiogenesis, the scope of BfBio extends beyond this specific biological process. As long as relevant co expression and PPI datasets and suitable training genes are available, this tool could be applied to any specific cell type, disease or condition to prioritize genes of interest.

## Supporting information

**S1 Appendix. Supplementary Figures.**

**S1 Table Set of 50 characterized EC stalk-cell genes used as training genes in this work.**

**S2 Table Top 49 ranked genes in the integrated network with the PPR algorithm.**

**S3 Table Top 49 genes prioritized by ToppNet.**

**S4 Table Top 49 genes prioritized by the original PageRank algorithm.**

**S5 Table PubMed Search results for the top 49 ranked genes with our PPR algorithm.**

**S6 Table All available studies from DiSign Atlas related to the 4 candidate genes.**

## Acknowledgments

We thank Mieke Dewerchin, Franky Terras, Jakub Salagovic and Jeffrey Kroon for their insightful comments.

## Data and Software Availability

The codes and data of BfBio are available on GitHub at github.com/AngioChange/Brain-for-Biotech.

