## Supplementary figures for "BfBio: a graph-based tool for the prediction of Angiogenic Stalk Cell genes using a Personalized PageRank algorithm"

**Fig S1. Enrichment Analysis for the 20 genes prioritized with GeneMANIA.**

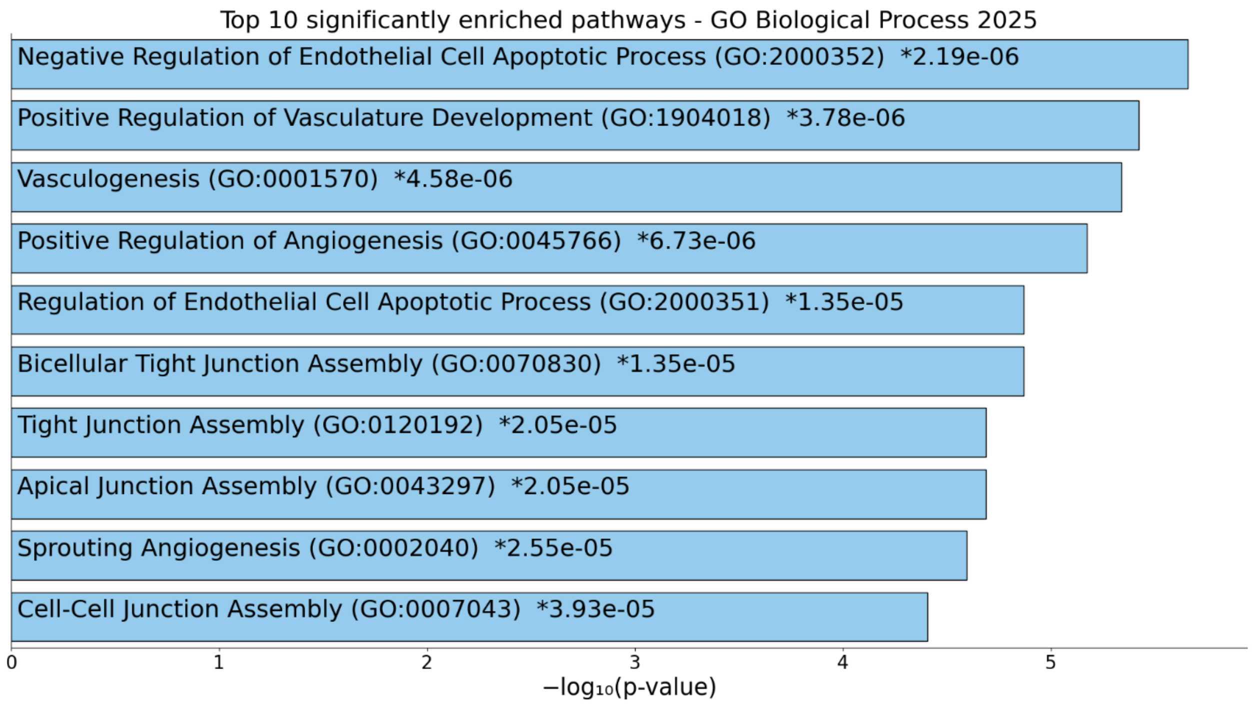

**Fig S2. Enrichment Analysis for the 49 genes prioritized with ToppNet.**

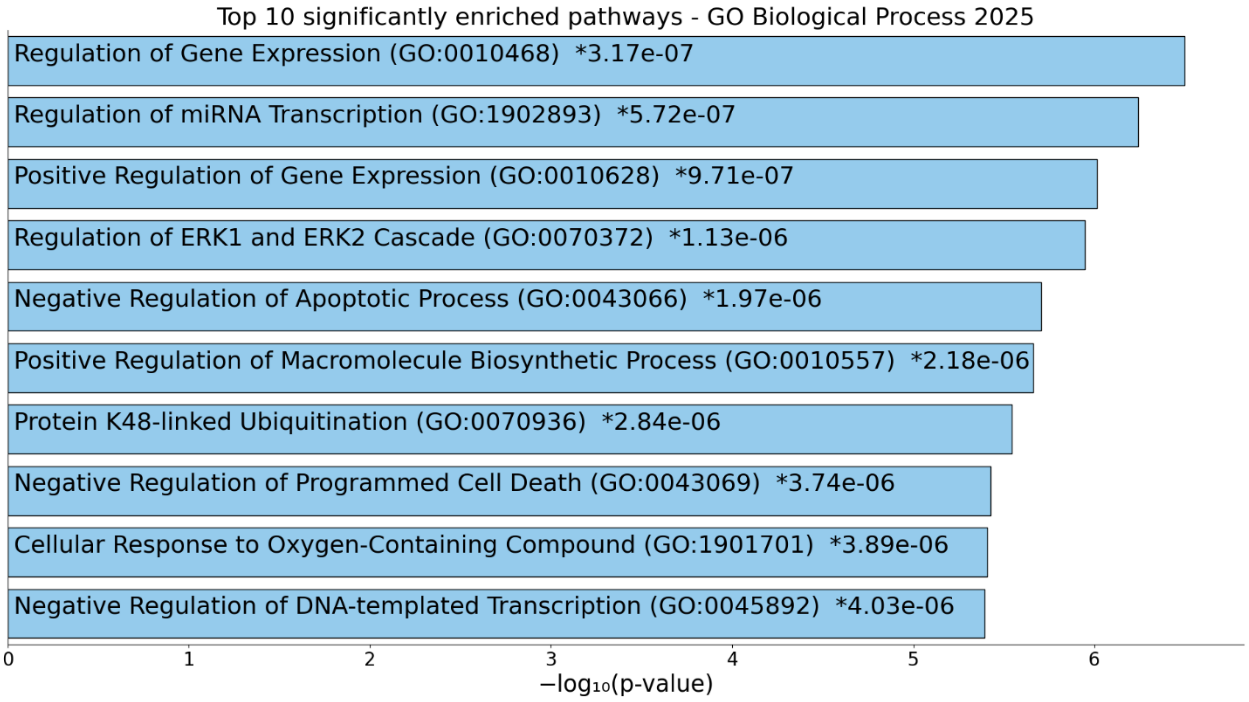

**Fig S3. Enrichment Analysis for the 49 genes prioritized with the original PageRank algorithm.**

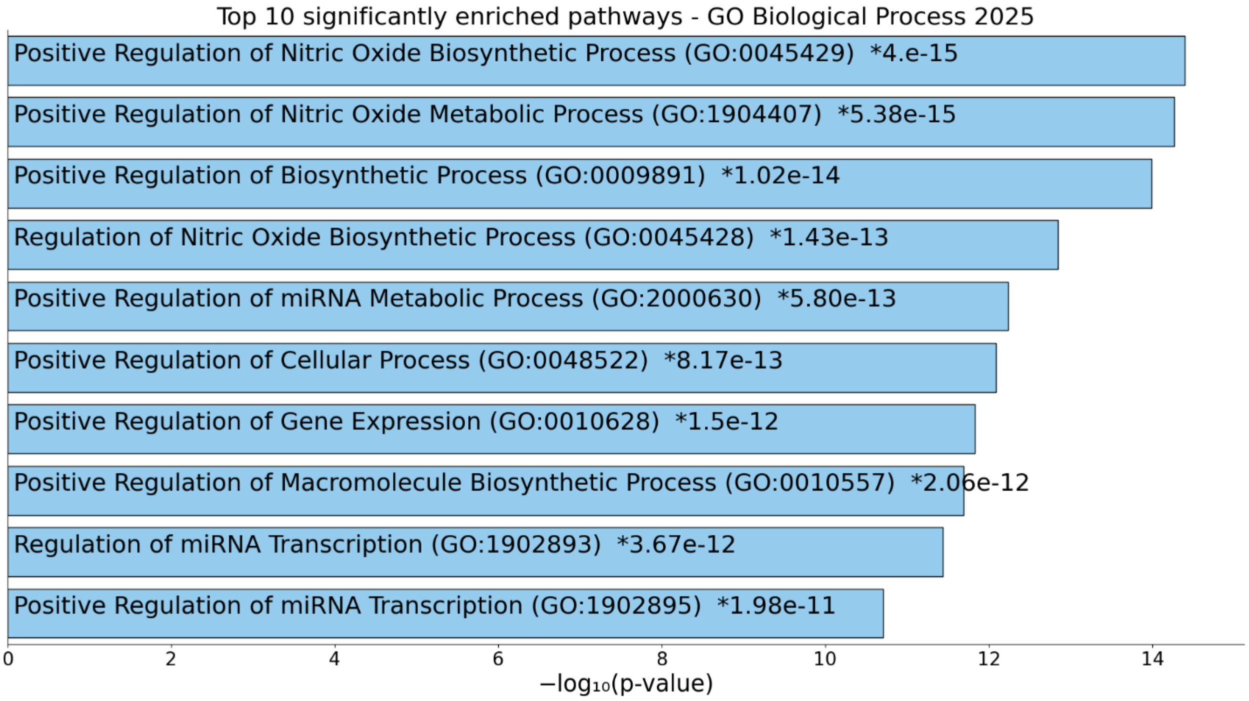

**Fig S4. AlphaFold-predicted 3D structures of the 4 potential target genes.**

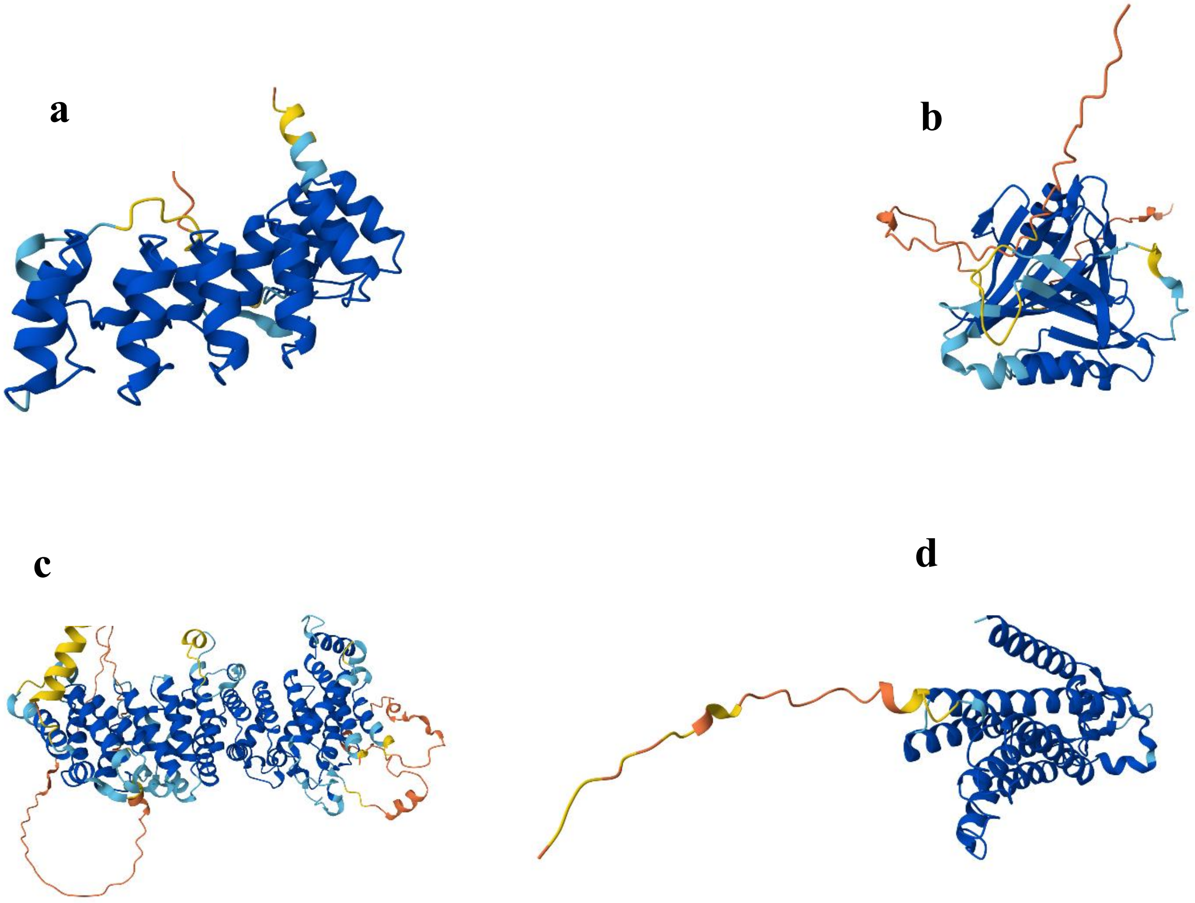


**a**. 3D structure of *ANKRD36B*. **b.** 3D structure of *FAM124A*. **c.** 3D structure of *FHIP2A*. **d.** 3D structure of *PLPP4*.

**Fig S5. Predicted topology visualization of *PLPP4* from Protter identifying 6 transmembrane domains.**

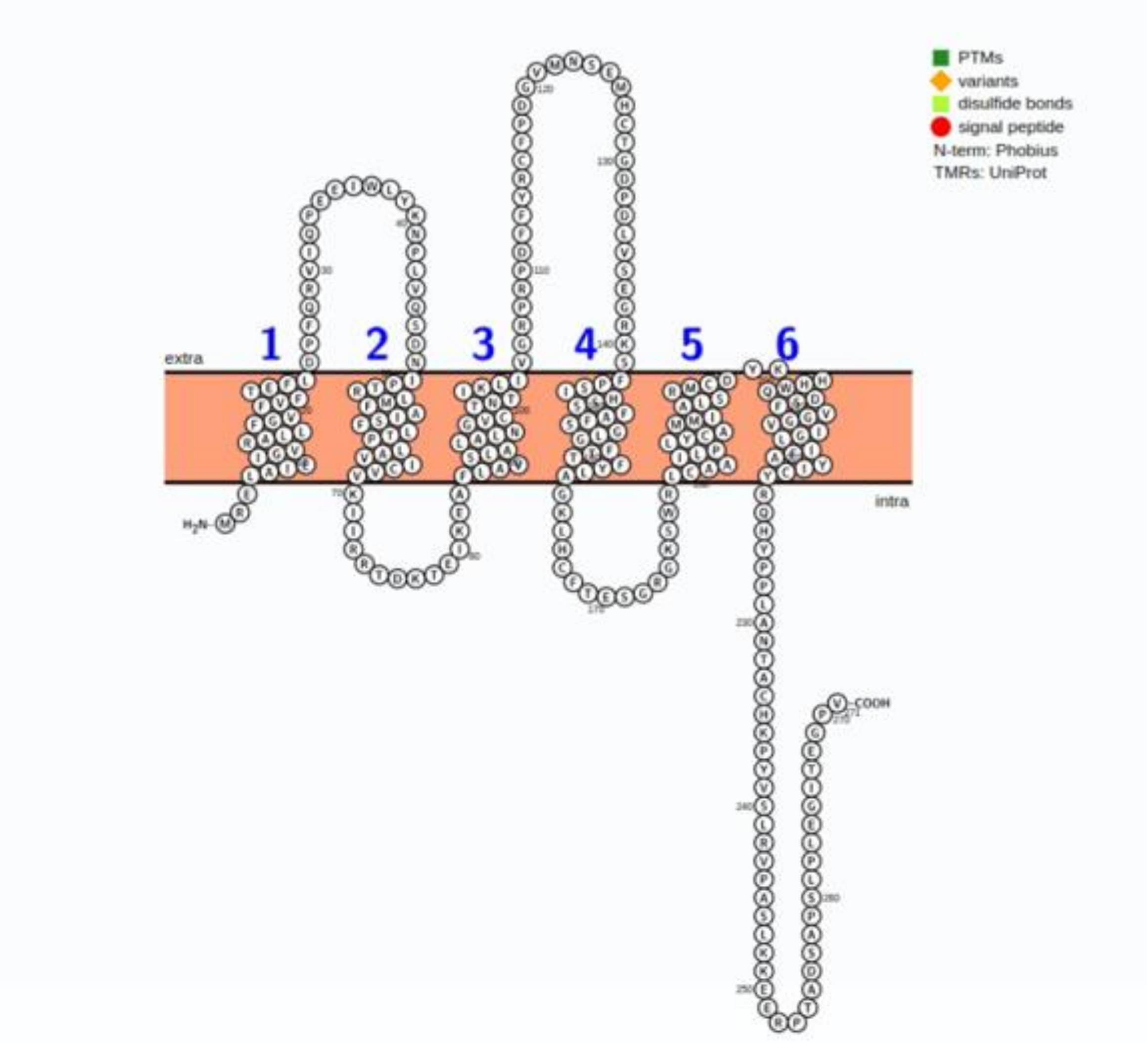
